# Non-replication of functional connectivity differences in autism spectrum disorder across multiple sites and denoising strategies

**DOI:** 10.1101/640797

**Authors:** Ye He, Lisa Byrge, Daniel P Kennedy

## Abstract

A rapidly growing number of studies on autism spectrum disorder (ASD) have used resting-state fMRI to identify alterations of functional connectivity, with the hope of identifying clinical biomarkers or underlying neural mechanisms. However, results have been largely inconsistent across studies, and there is therefore a pressing need to determine the primary factors influencing replicability. Here, we used resting-state fMRI data from the Autism Brain Imaging Data Exchange to investigate two potential factors: denoising strategy and data site (which differ in terms of sample, data acquisition, etc.). We examined the similarity of both group-average functional connectomes and group-level differences (ASD vs. control) across 33 denoising pipelines and four independently-acquired datasets. The group-average connectomes were highly consistent across pipelines (r = 0.92±0.06) and sites (r = 0.88±0.02). However, the group differences, while still consistent within site across pipelines (r = 0.76±0.12), were highly inconsistent across sites regardless of choice of denoising strategies (r = 0.07±0.04), suggesting lack of replication may be strongly influenced by site and/or cohort differences. Across-site similarity remained low even when considering the data at a large-scale network level or when considering only the most significant edges. We further show through an extensive literature survey that the parameters chosen in the current study (i.e., sample size, age range, preprocessing methods) are quite representative of the published literature. These results highlight the importance of examining replicability in future studies of ASD, and, more generally, call for extra caution when interpreting alterations in functional connectivity across groups of individuals.

## Introduction

Autism spectrum disorder (ASD) is a neurodevelopmental disorder with heterogeneous etiology and phenotypic expression. Resting-state functional Magnetic Resonance Imaging (rs-fMRI) --in which the temporal coupling of spontaneous activity across the brain, or functional connectivity (FC; Biswal, Yetkin, Haughton, & Hyde, 1995; Greicius, Krasnow, Reiss, & Menon, 2003), is measured -- has been widely used to study differences in functional brain organization in ASD, with hopes of revealing underlying neural mechanisms or identifying FC-based biomarkers (Abraham et al., 2017; Yahata et al., 2016). However, findings of FC alterations in ASD have been highly variable across studies (Hull et al., 2016). This variability of findings may reflect the variability across numerous study-specific factors, including strategies for denoising the data (i.e., preprocessing pipelines) and a host of differences across sites. Yet, without replicable findings that generalize beyond a single study, the utility of rs-fMRI for identifying mechanisms or serving as biomarkers of ASD is uncertain and remains to be demonstrated.

One potential source of variability across rs-fMRI studies has been the methods used for data preprocessing. The blood oxygenation-level dependent (BOLD) signal, while sensitive to changes related to brain activity, is also highly vulnerable to head motion and physiological noise, which can spuriously influence measures of functional connectivity and ultimately affect conclusions from functional connectivity studies (Power, Barnes, Snyder, Schlaggar, & Petersen, 2012; Power et al., 2014; Satterthwaite et al., 2012; Van Dijk, Sabuncu, & Buckner, 2012; Yan, Cheung, et al., 2013; Dadi et al., 2019). Ideally, effective data preprocessing methods would minimize the influence of such nuisance signals and improve reproducibility. Best practices for denoising methods are still evolving and a consensus has yet to be reached, in part because our understanding of how such artifacts influence the BOLD signal remains incomplete (Birn, 2012; Byrge & Kennedy, 2018; Power, Plitt, Laumann, & Martin, 2017). These differences presumably contribute in part to inconsistencies across studies -- different strategies have been used both within and across labs, adding additional uncontrolled and unaccounted for variation in the research literature. Even when researchers attempt to conduct *post hoc* analyses to try to understand how different preprocessing steps could account for study-level differences in ASD, the lack of a ground truth upon which to evaluate measurement accuracy limits our ability to interpret such differences (Müller et al., 2011).

Most common denoising approaches rely on linear regression, whereby various estimates of noise are regressed from the BOLD data. The numerous variations of this strategy come from different choices of which noise estimates to use as regressors. Those most commonly used include measures of head displacement along six translational and rotational dimensions, as well as time series from white matter (WM) and cerebrospinal fluid (CSF). An especially controversial nuisance regressor is the global fMRI signal; proponents of global signal regression (GSR) argue for its efficacy in removing physiological noise (Birn, 2012; Byrge & Kennedy, 2018; Power et al., 2017), while the concerns include removal of real neural signals (Scholvinck, Maier, Ye, Duyn, & Leopold, 2010) and distorting clinical group comparison (Gotts et al., 2013; Yang et al., 2014). An additional preprocessing step that can be used in parallel is volume censoring (or “scrubbing”; Power et al., 2012), in which specific time points associated with excessive amounts of framewise displacement (FD; corresponding to moments of head movement) and/or changes in global signal are excluded from analysis. A related choice is called “spike regression”, which regresses from the data one or more nuisance regressors labeling time points contaminated with excessive motion (Lemieux, Salek-Haddadi, Lund, Laufs, & Carmichael, 2007; Satterthwaite et al., 2013).

Several recent studies have evaluated the performance of different denoising strategies. Although no relationship between motion and functional connectivity should remain following an optimal denoising procedure, these studies found that the strength of residual relationships between FC and artifacts varied widely across commonly-used pipelines (Byrge & Kennedy, 2018; Ciric et al., 2017; Parkes, Fulcher, Yucel, & Fornito, 2018). Given that greater in-scanner head movement is commonly observed in ASD and other clinical populations, differences in preprocessing choices and particularly how those choices deal with artifacts arising from head movement could be a potential source of inconsistent results across rs-fMRI studies. For example, Gotts and colleagues (2013) compared the effects of pipelines with and without GSR on group comparisons of functional connectivity between ASD and controls. They found that group differences varied across pipelines and demonstrated that GSR affected group comparison results. Jones et al. (2010) also found that the use of GSR influenced findings of group differences in connectivity in ASD. Parker and colleagues (2018) systematically evaluated the influence of numerous denoising pipelines on group differences in functional connectivity in schizophrenia. They found that significant group differences were only found in some pipelines (including GSR and aCompCor) and that the overlap between functional connections (i.e., edges) identified in different pipelines was generally low. These findings demonstrate clearly that the choice of denoising pipeline can affect the results of clinical comparisons, including both the presence or absence of group differences and their specific details (e.g., specific edges affected).

Further complicating the picture is that data site effects, or variation across different scanning sites, have been reported in several studies of both task-based and resting-state fMRI (Brown et al., 2011; Dansereau et al., 2017; Noble et al., 2017; Turner et al., 2013; Yamashita et al., 2019; Yan, Craddock, Zuo, Zang, & Milham, 2013; Yu et al., 2018). Different data sites present many potential sources of variation, including differences in participant (i.e., cohort) characteristics, image acquisition parameters, scanners, scan procedures, and more. Such uncontrolled variation could undermine the generalizability of results and efforts to uncover underlying mechanisms and clinically useful biomarkers. Clinical and etiological heterogeneity within the ASD population could also exacerbate these difficulties. Nair et al. (2018) compared a local measure of functional connectivity (ReHo, regional homogeneity) between ASD and controls from different samples. They found few consistent results across samples, even when using the same analysis pipeline and examining only data collected with eyes open. They suggested that extra caution should be paid to between-site variability when using multi-site data. King et al. (2019) examined many FC-related measures on group differences between ASD and the control. They found none of the measures could wholly reproduce the group differences across sites. However, a recent study reported reproducible ASD-associated alterations of functional connectivity across four large ASD cohorts (Holiga et al., 2019).

The Autism Brain Imaging Data Exchange, or ABIDE, provides the ideal data in which to test the influence of some of these factors. ABIDE is a data sharing initiative wherein researchers across laboratories shared resting-state data from TD and ASD participants for the flexible use by other researchers, “allow[ing] for replication, secondary analyses and discovery efforts” (Di Martino et al., 2014). This flexibility has allowed a proliferation of research on functional connectivity in ASD, and the data has been used in various ways, including considering each site separately (Hahamy, Behrmann, & Malach, 2015; Pua, Malpas, Bowden, & Seal, 2018) or using multi-site aggregation (Abraham et al., 2017; Floris, Lai, Nath, Milham, & Di Martino, 2018; King et al., 2019). Both of these approaches are widely employed, and while greater statistical power can be achieved from aggregation, examining multiple individual sites can be used to evaluate replicability. Here, we used four of the largest ABIDE sites to quantify 1) the similarity/variation of ASD-control group differences of FC across denoising pipelines, 2) the similarity/variation of ASD-control group differences across data sites, 3) and the effect of pipelines on inter-site variation.

## Methods

### Literature survey

A literature survey was conducted to summarize the usage of denoising methods and sample characteristics (age range and sample size etc.) from recent resting-state FC-MRI studies of ASD. We searched the PubMed database using keywords consisted of “resting*”, “autism”, and “fMRI”, or combining “resting*”, “autism” and “connectivity”, published from the beginning of 2013 until June 2019 (inclusive of the time when ABIDE data has been available). In total, 245 studies were identified. To be in line with our study focusing on case-control comparison of resting-state functional connectivity, 118 studies were excluded for either not using fMRI, using animals, not analyzing static functional connectivity, no group comparison, focusing on machine learning to classify, reviews or not accessible.

### Participants

To enable an accurate evaluation of factors affecting replication of ASD-related FC alterations obtained by typical study design, four independent data sites (NYU, SDSU, UCLA, and UM) from ABIDE I and ABIDE II were analyzed (Di Martino et al., 2017; Di Martino et al., 2014). We chose these four sites in consideration of their large sample sizes as well as maximally overlapping age ranges of participants across all four sites (Table 1 and Figure S1). For example, although USM also has large sample size, the number of participants whose ages overlapped with other sites is limited; therefore, we did not include this site in our analysis. To reduce variability while maximizing sample size, we included participants based on following criteria: 1) age ranging from 10 to 20 years old; 2) IQ > 70; 3) mean FD no larger than 0.3 mm; 4) sufficient quality of anatomical images, assessed by manual checking. To further control potential head-motion differences between groups, we matched each single ASD participant with a control participant with the smallest difference in mean FD within each site, and removed any additional subjects not matched. ASD and typically developing (TD) control participants were not significantly different on mean FD or mean translation or rotation movement parameters for any site (Table 1).

**Table 1.** Demographic information

|  | NYU |  |  | SDSU |  |  | UCLA |  |  | UM |  |  |
| --- | --- | --- | --- | --- | --- | --- | --- | --- | --- | --- | --- | --- |
|  | ASD | TD | Stats<br>(p/t) | ASD | TD | Stats<br>(p/t) | ASD | TD | Stats<br>(p/t) | ASD | TD | Stats<br>(p/t) |
| n | 44 | 44 | - | 36 | 36 | - | 38 | 38 | - | 34 | 34 | - |
| Male/<br>Female | 38/6 | 38/6 | - | 32/4 | 29/7 | - | 37/1 | 31/7 | - | 28/6 | 28/6 | - |
| Age (mean±<br>SD) | 13.0 ±<br>2.5 | 13.7 ±<br>2.4 | 0.24/<br>-1.2 | 14.3±<br>2.5 | 14.2 ±<br>2.1 | 0.72/<br>0.4 | 13.6 ±<br>2.3 | 13.3 ±<br>1.6 | 0.63/<br>0.49 | 14.6 ±<br>1.9 | 14.7 ±<br>2.5 | 0.78/<br>-0.3 |
| IQ<br>(mean±SD) | 102.0<br>± 13.6 | 113.1<br>± 15.0 | *0.00/<br>-3.7 | 104.7<br>± 16.1 | 104.2<br>± 11.2 | 0.87/<br>0.2 | 103.4<br>± 13.0 | 106.8<br>± 10.5 | 0.21/<br>-1.3 | 109.3<br>± 15.8 | 109.6<br>± 10.4 | 0.93/<br>-1.0 |
| Handedness<br>(R/L/A) | - | - | - | 29/5/2 | 31/3/2 | - | 33/5/0 | 35/3/0 | - | 25/5/0 | 29/4/0 | - |
| ADI social<br>(mean± SD, n) | 19.9 ±<br>6.2<br>(41) | - | - | 18.3 ±<br>4.2<br>(34) | - | - | 20.8 ±<br>4.6<br>(28) | - | - | 19.5 ±<br>4.7<br>(34) | - | - |
| ADI verbal<br>(mean± SD, n) | 15.7 ±<br>4.7<br>(42) | - | - | 13.1 ±<br>4.8<br>(34) | - | - | 16.6 ±<br>4.4<br>(28) | - | - | 15.8 ±<br>3.2<br>(34) | - | - |
| ADI RRB<br>(mean± SD, n) | 5.8 ±<br>2.8<br>(42) | - | - | 5.8 ±<br>2.3<br>(34) | - | - | 7.4 ±<br>2.0<br>(28) | - | - | 5.9 ±<br>2.5<br>(34) | - | - |
| ADOS total<br>(mean± SD, n) | 11.6 ±<br>4.0<br>(44) | - | - | 13.5 ±<br>5.0<br>(35) | - | - | 11.6 ±<br>3.8<br>(36) | - | - | 11.3 ±<br>4.6<br>(41) | - | - |
| SRS (mean±<br>SD, n) | 91.3 ±<br>32.8<br>(43) | 24.2 ±<br>13.3<br>(39) | 0.07/<br>1.8 | 101.7<br>± 22.8<br>(24) | 21.1 ±<br>10.9<br>(18) | 0.00/<br>13.8 | - | - | - | - | - | - |
| N medicated<br>(n reporting) | 10(42) | 0(44) | - | 13(36) | 0(36) | - | 20(38) | 1(38) | - | 15(34) | 1(33) | - |
| Mean FD<br>(mean± SD) | 0.15 ±<br>0.05 | 0.14 ±<br>0.04 | 0.32/<br>1.0 | 0.12 ±<br>0.06 | 0.12 ±<br>0.06 | 0.90/<br>1.0 | 0.16 ±<br>0.06 | 0.14 ±<br>0.07 | 0.18/<br>1.4 | 0.17 ±<br>0.06 | 0.15 ±<br>0.05 | 0.41/<br>0.8 |
| Mean<br>Translation<br>(mean± SD) | 0.16 ±<br>0.09 | 0.19 ±<br>0.15 | 0.26/-<br>1.1 | 0.13 ±<br>0.07 | 0.14 ±<br>0.10 | 0.69/-<br>0.4 | 0.15 ±<br>0.08 | 0.14 ±<br>0.09 | 0.71/0<br>.4 | 0.27 ±<br>0.19 | 0.22 ±<br>0.11 | 0.20/1<br>.3 |
| Mean<br>Rotation<br>(mean± SD) | 0.003<br>±<br>0.002 | 0.004<br>±<br>0.002 | 0.20/-<br>1.3 | 0.003<br>±<br>0.002 | 0.003<br>±<br>0.002 | 0.66/-<br>0.4 | 0.003<br>±<br>0.001 | 0.003<br>±<br>0.002 | 0.76/0<br>.3 | 0.005<br>±<br>0.004 | 0.004<br>±<br>0.003 | 0.60/0<br>.5 |
| Scrubbed<br>Volumes<br>(mean± SD) | 36.3±<br>23.8 | 34.5±<br>23.5 | 0.73/<br>0.4 | 25.7±<br>20.5 | 24.4±<br>24.8 | 0.82/<br>0.2 | 29.1 ±<br>17.5 | 22.9 ±<br>16.5 | 0.12/<br>1.6 | 66.9±<br>47.1 | 59.6±<br>36.5 | 0.48/<br>0.7 |

### Image Preprocessing

The rs-fMRI scanning parameters for each site are shown in Table 2. All the images were preprocessed using Matlab (R2018a) code made available from a recent study (Parkes et al., 2018) that integrates SPM 12, FSL (FMRIB’s Software Library; Smith et al., 2004) and Advanced Normalization Tools (ANTs; Avants, Epstein, Grossman, & Gee, 2008). The T1 images were preprocessed using the following steps: neck removal; segmentation of white matter (WM), cerebral spinal fluid (CSF), and grey matter (GM); five times erosion of WM mask and two times erosion of CSF mask; nonlinear registration of T1 images to MNI space, and applying the transformation to WM, CSF, and GM masks.

**Table 2.** rs-fMRI scanning parameters. Note that there are several parameters that are different across these sites. The current study was not meant to control specifically for each of these (e.g., UCLA has a 3000 msec TR vs. all other sites with 2000 msec) because such differences are also present in published studies where such factors are not controlled, but where replication is still implicitly expected. These datasets are also sometimes aggregated together, again implicitly assuming that such differences will not have major effects on case-control differences. Note also that the UM site acquired more volumes per participant than the other sites; we chose not to downsample this data for our main anaylysis because including more data from each individual participant should yield a better estimate of an individual’s functional connectivity; in other words, downsampling would produce artificially noisier data, would not be representative of the actual data available and analyzed in other published reports, and would bias our results away from finding evidence for across-site replication. However, we did rerun the primary analyses using a downsampled version of the UM data and findings remained the same.

|  | NYU | SDSU | UCLA | UM |
| --- | --- | --- | --- | --- |
| Scanner | Siemens 3T Allegra | GE 3T MR750 | Siemens 3T TIM Trio | GE 3T Signa |
| TR/TE | 2000/15 | 2000/30 | 3000/28 | 2000/30 |
| FA | 90 | 90 | 90 | 90 |
| Resolution | 3×3×4 | 3.4×3.4×3.4 | 3×3×4 | 3.4×3.4×3 |
| Volumes | 180 | 180 | 120 | 300 |
| Matrix | 64×80×33 | 64×64×42 | 64×64×34 | 64×64×40 |

Preprocessing of functional images included several steps shared across different denoising pipelines, including the following: removing the first four volumes; slice-timing correction; head motion correction by volume realignment; co-registration to the native structural image using rigid-body registration, and then to the MNI template using nonlinear transformations derived from T1 registration; removing linear trends; normalization of global mean intensity to 1000 units; conducting different denoising strategies (detailed in the next section); bandpass filtering (0.008 −0.08 Hz); and spatial smoothing with a 6 mm full-width at half-maximum filter.

### Denoising Pipelines

We analyzed imaging data using several commonly-used denoising methods, together with various combinations of different nuisance regressors and volume censoring approaches, resulting in a total of 33 denoising pipelines (Table 3).

**Table 3.** Compositions of Denoising Pipelines

| Denoising Pipelines | Head motion parameters | Tissue-based Regressors | GSR | Censoring |
| --- | --- | --- | --- | --- |
| 6H | 6 | - | - | - |
| 12H | 12 | - | - | - |
| 24H | 24 | - | - | - |
| 6H+2W | 6 | mean WM/CSF | - | - |
| 12H+2W | 12 | mean WM/CSF | - | - |
| 24H+2W | 24 | mean WM/CSF | - | - |
| 24H+4W | 24 | 4 mean WM/CSF | - | - |
| 24H+8W | 24 | 8 mean WM/CSF | - | - |
| 6H+aCC | 6 | aCompCor | - | - |
| 12H+aCC | 12 | aCompCor | - | - |
| 24H+aCC | 24 | aCompCor | - | - |
| 6H+2W+Spike | 6 | mean WM/CSF | - | Spike |
| 6H+2W+Scrub | 6 | mean WM/CSF | - | Scrub |
| 12H+2W+Spike | 12 | mean WM/CSF | - | Spike |
| 12H+2W+Scrub | 12 | mean WM/CSF | - | Scrub |
| 24H+2W+Spike | 24 | mean WM/CSF | - | Spike |
| 24H+2W+Scrub | 24 | mean WM/CSF | - | Scrub |
| 6H+2W+GSR | 6 | mean WM/CSF | 1 | - |
| 12H+2W+GSR | 12 | mean WM/CSF | 1 | - |
| 24H+2W+GSR | 24 | mean WM/CSF | 1 | - |
| 24H+4W+GSR | 24 | 4 mean WM/CSF | 1 | - |
| 24H+8W+4GSR | 24 | 8 mean WM/CSF | 4 | - |
| 6H+aCC+GSR | 6 | aCompCor | 1 | - |
| 12H+aCC+GSR | 12 | aCompCor | 1 | - |
| 24H+aCC+GSR | 24 | aCompCor | 1 | - |
| 6H+2W+GSR+Spike | 6 | mean WM/CSF | 1 | Spike |
| 6H+2W+GSR+Scrub | 6 | mean WM/CSF | 1 | Scrub |
| 12H+2W+GSR+Spike | 12 | mean WM/CSF | 1 | Spike |
| 12H+2W+GSR+Scrub | 12 | mean WM/CSF | 1 | Scrub |
| 24H+2W+GSR+Spike | 24 | mean WM/CSF | 1 | Spike |
| 24H+2W+GSR+Scrub | 24 | mean WM/CSF | 1 | Scrub |
| ICA-AROMA+2W | - | mean WM/CSF | - | - |
| ICA-AROMA+2W+GSR | - | mean WM/CSF | 1 | - |

#### Regression of head motion parameters

Head motion parameters are based on six time series reflecting in-scanner head movements along three translational axes and three rotational axes. We examined three variants: 6H (just these original 6 motion parameters), 12H (including the original 6H, plus the first derivative of each as computed by backward differences), and 24H (including 12H, plus the squares of each of the 12 parameters) (Satterthwaite et al., 2013).

#### Regression of signals from white matter and cerebrospinal fluid

We used two methods to estimate WM and CSF signals: (a) mean WM/CSF, the average time series across voxels within WM and CSF masks, with three variants: mean WM and CSF alone (2W), or adding their temporal derivatives (4W), or adding squares of 4W (8W), and (b) aCompCor, which applies principal component analysis to the time series from WM and CSF voxels separately, and uses the top five principal components for each tissue compartment (Muschelli et al., 2014).

#### Regression of global mean signal

Global mean signal was calculated by averaging voxel-wise time series across the whole brain (GSR) or extended with squares of it and their temporal derivatives (4GSR).

#### Volume Censoring

Volume censoring involves censoring specific time points in BOLD data that have excessive head motion, which was evaluated using framewise displacement (FD). We adopted two different censoring strategies: spike regression and scrubbing. To keep consistent with previous work, we calculated FD differently for spike regression and scrubbing and used different thresholds. For spike regression, FD was calculated as the root mean square of framewise changes of six head motion parameters (Jenkinson, Bannister, Brady, & Smith, 2002; Satterthwaite et al., 2013). This FD trace was then used as an additional nuisance regressor in which volumes with FD above 0.25 mm were marked as 1 and otherwise as 0, which was then regressed (together with other regressors) from the BOLD time series. For scrubbing, FD was calculated as the sum of absolute framewise changes of six head motion parameters (Power et al., 2012). Volumes with FD above 0.2mm were excluded from analysis at the end of preprocessing. We excluded subjects with less than 4 minutes of valid BOLD data following spike regression or scrubbing.

#### ICA-AROMA

ICA-AROMA uses independent component analysis (ICA) to decompose the BOLD signal into spatial independent components and then automatically identify motion-related components based on assessing high-frequency content, correlation with head realignment parameters, edge fraction and CSF fraction of each component (Pruim et al., 2015). ICA-AROMA is performed for each participant separately and the number of motion-related components can vary for different participants. Spatial smoothing was performed before noise regression when using ICA-AROMA.

### Functional Connectome Construction

We used a parcellation template containing 200 cortical ROIs to construct the functional connectome for each subject (Schaefer et al., 2018). Specifically, after preprocessing we weight-averaged the time series of all voxels within each ROI based on their grey matter probability. Then we computed the Pearson’s correlation between time series of each pair of 200 ROIs to construct a 200 by 200 functional connectivity matrix of each pipeline for each subject, and Fisher-z transformed correlation coefficients for the purpose of normalization. The group-average functional connectome was obtained by averaging functional connectomes across participants of each group for further analysis.

### Group Differences between ASD and Controls

We compared the ASD to the control group on each edge in the functional connectome matrix for each dataset, using the non-parametric Wilcoxon rank sum tests to reduce the influence of extreme data. Age and mean FD were first regressed out as covariates. A 200 × 200 statistic z-value map (z-map) representing group differences for all edges was obtained for each pipeline in each dataset.

### Assessing Replicability of Whole Functional Connectomes

A schematic is shown in Figure 1 to illustrate our approach. We first averaged functional connectomes across participants with ASD and across typical controls, separately for each pipeline and for each site. Next, to assess the similarity of functional connectomes across denoising methods, we calculated the Spearman’s correlation of group-average functional connectomes between each pair of pipelines to derive a similarity matrix, separately within each data site. To better visualize the distance between pipelines, we used multi-dimensional scaling (MDS) to transform each pipeline-similarity matrix into a representation in two-dimensional space. Each point corresponds to a different pipeline and the distance between points corresponds to their degree of dissimilarity. We used Procrustes analysis (without scaling) to best align the plots across sites, using NYU as the reference plot. To assess the across-site similarity of functional connectomes, we calculated the Spearman’s correlation between each pair of four datasets under each pipeline.

**Figure 1.**
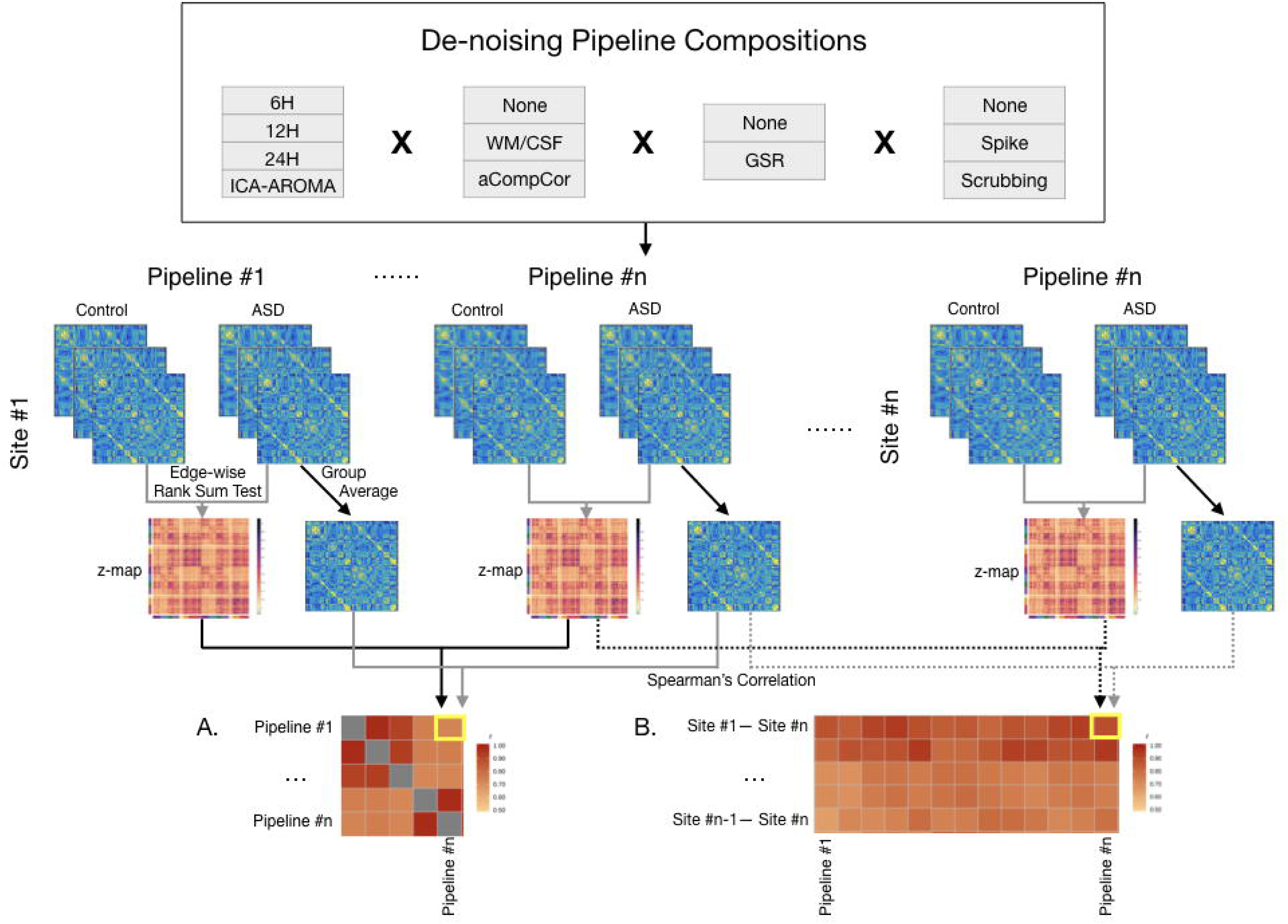
Schematic plot for post-processing analysis. We used a total of 33 denoising pipelines, with different combinations of regression of head motion parameters (6H/12H/24H), ICA-AROMA, signals of white matter/cerebral spinal fluid (WM/CSF), global mean signal (GSR), and volume censoring (spike/scrubbing). Functional connectomes were separately constructed with 33 pipelines for each subject. We averaged functional connectomes across each subject group as well as compared each cell in the connectome between two groups to derive group-difference z-maps. Then we calculated the Spearman’s correlation between group-average functional connectomes, as well as between z-maps (a) across pipelines and (b) across datasets.

### Assessing Replicability of Group Differences between ASD and Controls

To evaluate the similarity of ASD-control group differences across denoising methods or sites, we calculated the Spearman’s correlation between whole brain group difference (z-map) matrices across pipelines within each site, as well as across sites.

In addition to comparing whole brain z-map matrices, we further focused on those edges showing the greatest difference between the ASD and control groups for across sites comparisons. First, we sort all edges based on their z values for each pipeline in each data site, and obtained top 500 (positive, ASD > control) and bottom 500 (negative, ASD < control) edges (approximately 5% of total edges) for each map. Then we calculated how many those edges overlapped between each two maps (pipeline/site) separately for positive and negative z values. Permutation tests were used to examine whether the numbers of overlapping edges were above chance. First, we shuffled the diagnostic labels (ASD/control) of all the subjects within each site, keeping original sample sizes for each group. Then we compared these two new groups to derive a null z-map for each pipeline within each site, and then calculated the overlapping edges between sites using the same method as above. This procedure was repeated 1000 times for each pipeline to generate a null distribution of chance levels of overlapping edges across sites. So as to not be overly conservative, results are not corrected across the 33 pipelines examined, but are corrected for the six pairwise site comparisons (e.g., FDR correction, *q* <= .05; Benjamini & Hochberg, 1995).

We also examined the similarity of group differences between data sites at a large-scale network level. We mapped the whole functional connectome to a 17 functional networks template (Yeo et al., 2011) and obtained a 17 × 17 connectivity matrix by averaging connectivity of edges in each cell. Using the same statistical method to compare each cell between the ASD and control groups, we obtained a z-map for each pipeline in each site and calculated Spearman’s correlation across sites under each pipeline as described above.

### Data and code availability

All data is available from the ABIDE repository, preprocessing code was available from Parkes et al. (2018), and our code is available upon reasonable request.

## Results

### Literature survey

We sought to provide descriptive information regarding common preprocessing approaches of case-control studies of ASD, with the intent of contextualizing the parameters of the current study relative to the published literature. Figure 2 demonstrates the highly varied preprocessing methodologies applied in previous studies. Regression of head motion parameters is an extremely common step but varies in terms of its precise implementation (relatively equally split across 6, 12, or 24 parameters). Regression of average CSF and white matter signals is used more often than aCompCor (~55% vs ~28%). Just over half of studies used scrubbing (~49%) or spike regression (~8%) to remove the effects of motion-outlier volumes. Less than one third of studies used GSR (28%). ICA-AROMA is a recently developed method, and as such has only been used by a few studies to date (~4%).

**Figure 2.**
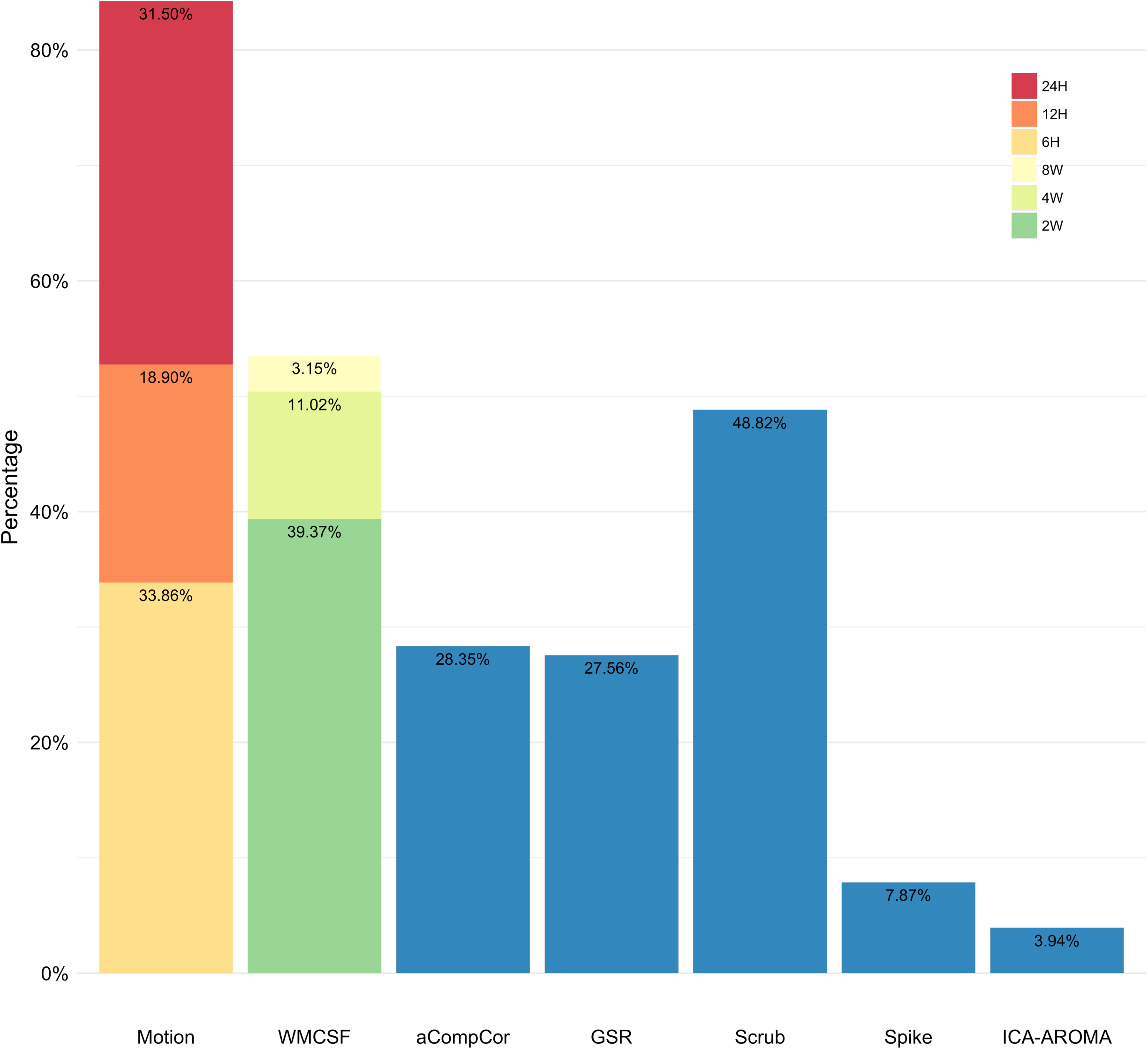
Usage proportion of different denoising preprocessing strategies in previous resting-state fMRI case-control studies on functional connectivity in ASD.

### Group-average functional connectome replicated across pipelines and across sites

We first assessed the similarity of group-average functional connectomes of ASD group across denoising pipelines separately within each data site. Generally, functional connectomes were highly similar across pipelines and this similarity pattern is consistent across sites (Figure 3A; NYU, *r* = 0.92±0.06; SDSU, *r* = 0.92±0.06; UCLA, *r* = 0.92±0.06; UM, *r* = 0.90±0.08). Results were similar for the control group (NYU, *r* = 0.91±0.07; SDSU, *r* = 0.93±0.06; UCLA, *r* = 0.93±0.05; UM, *r* = 0.90±0.08). As is apparent from the quadrant structure of Figure 3, GSR was a major influence on similarity of average functional connectomes across pipelines, such that similarity was extremely high with the same GSR status but reduced when pipelines differed in their use of GSR. We used multi-dimensional scaling to represent this graphically (Figure 3B), which demonstrates that the use of GSR is a primary dimension upon which results are either similar or different from one another.

**Figure 3.**
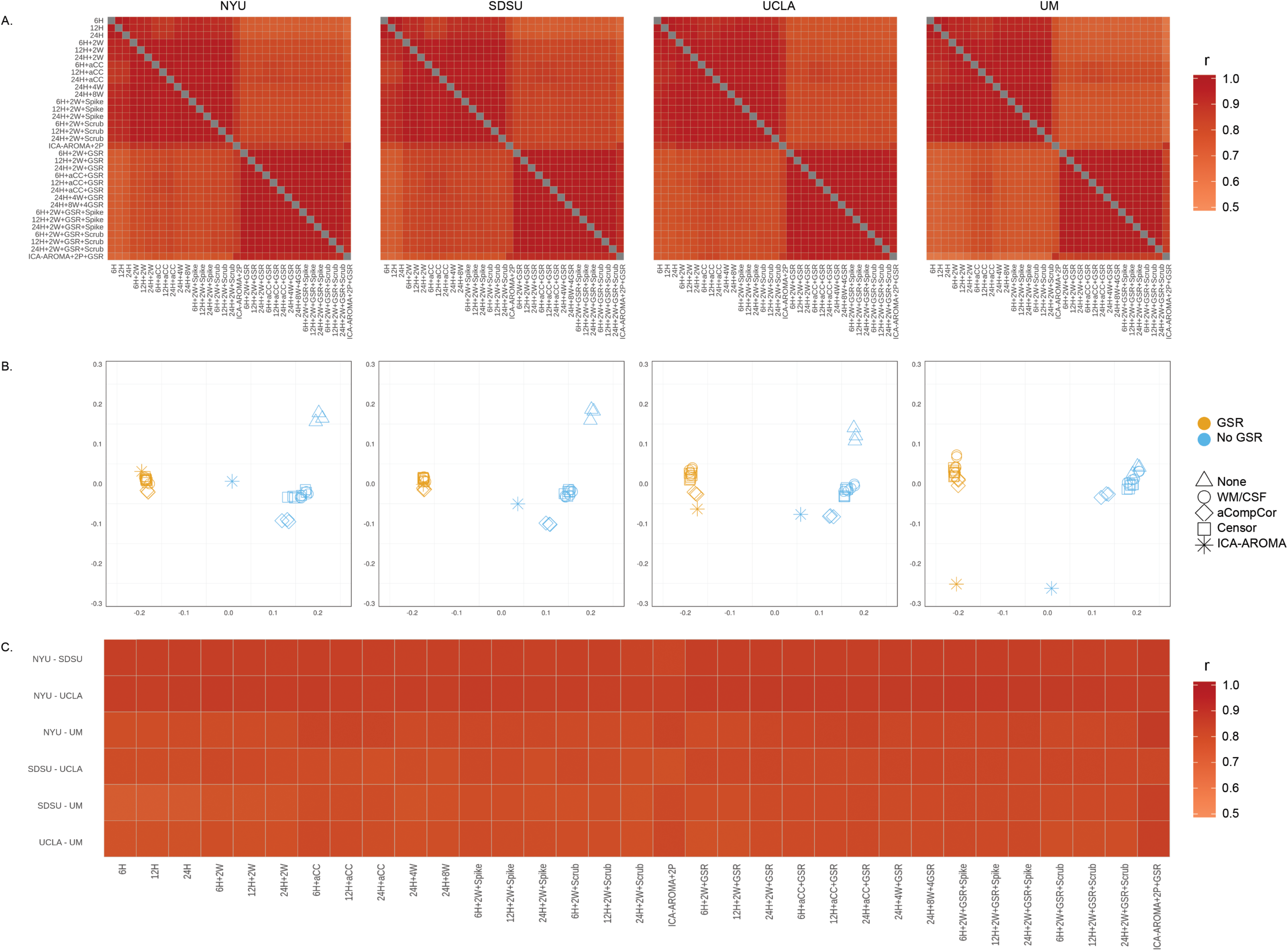
Consistency of group-average functional connectome across pipelines and sites. (A) Spearman’s correlation coefficients of group-average functional connectomes across pipelines. It indicates high similarity across pipelines, though pipelines with different GSR status were less similar, as is seen in quadrant structure. (B) It provides a different visualization of relative distance among different pipelines based on multi-dimensional scaling. Each data point represents a pipeline (note that not all points are visible because there is a high degree of overlap between some of them). It directly shows the major factor differentiating pipelines is based on the usage of GSR. The triangle shape corresponds to the basic pipelines (which only regress out 6H/12H/24H), the circle shape corresponds to the pipelines adding WM/CSF regression, the diamond shape corresponds to pipelines using aCompCor, the square shape corresponds to volume censoring (scrubbing and spiking) and the asterisk corresponds to ICA-AROMA. (C) Spearman’s correlation coefficients between group-average connectomes were inconsistently high across sites for all pipelines.

We examined the replicability of functional connectomes across sites under each pipeline. Figure 3C shows that group-average connectome of ASD is also similar across data sites within each pipeline (for all pipelines, *r* = 0.88±0.02). Pipelines with GSR increased between-site similarity compared to pipelines without GSR (rank sum, *z* = 3.88, *p* = 0.001). Note that first three minimally-preprocessed pipelines were excluded for this analysis because these tended to be quite different from all other approaches (as seen in Figure 3B).

In summary, group averaged functional connectomes were similar across pipelines and could be replicated across data sites – thus, even using different scanners and scanning protocols did not affect replicability of the group averaged functional connectome.

### Group-differences replicated across pipelines but not across sites

Next, we assessed similarity of ASD-control comparisons of functional connectomes across pipelines, within each site. The results were consistent across pipelines within each site for all four sites (NYU, *r* = 0.78±0.10; SDSU, *r* = 0.78±0.10; UCLA, *r* = 0.75±0.10; UM, *r* = 0.71±0.14). As in Figure 3, Figure 4 shows that GSR was also a dominant factor in similarity of group differences across pipelines – highly similar results with concordant use or non-use of GSR (i.e., either both present or absent), but reduced similarity when pipelines were discordant in their use of GSR (concordant: *r* = 0.84±0.09; discordant: *r* = 0.70±0.07, averaged across four sites), suggesting that the same data analyzed with or without GSR may yield different results. In addition, pipelines with ICA-AROMA were generally different from others (Figure 4 A and B).

**Figure 4.**
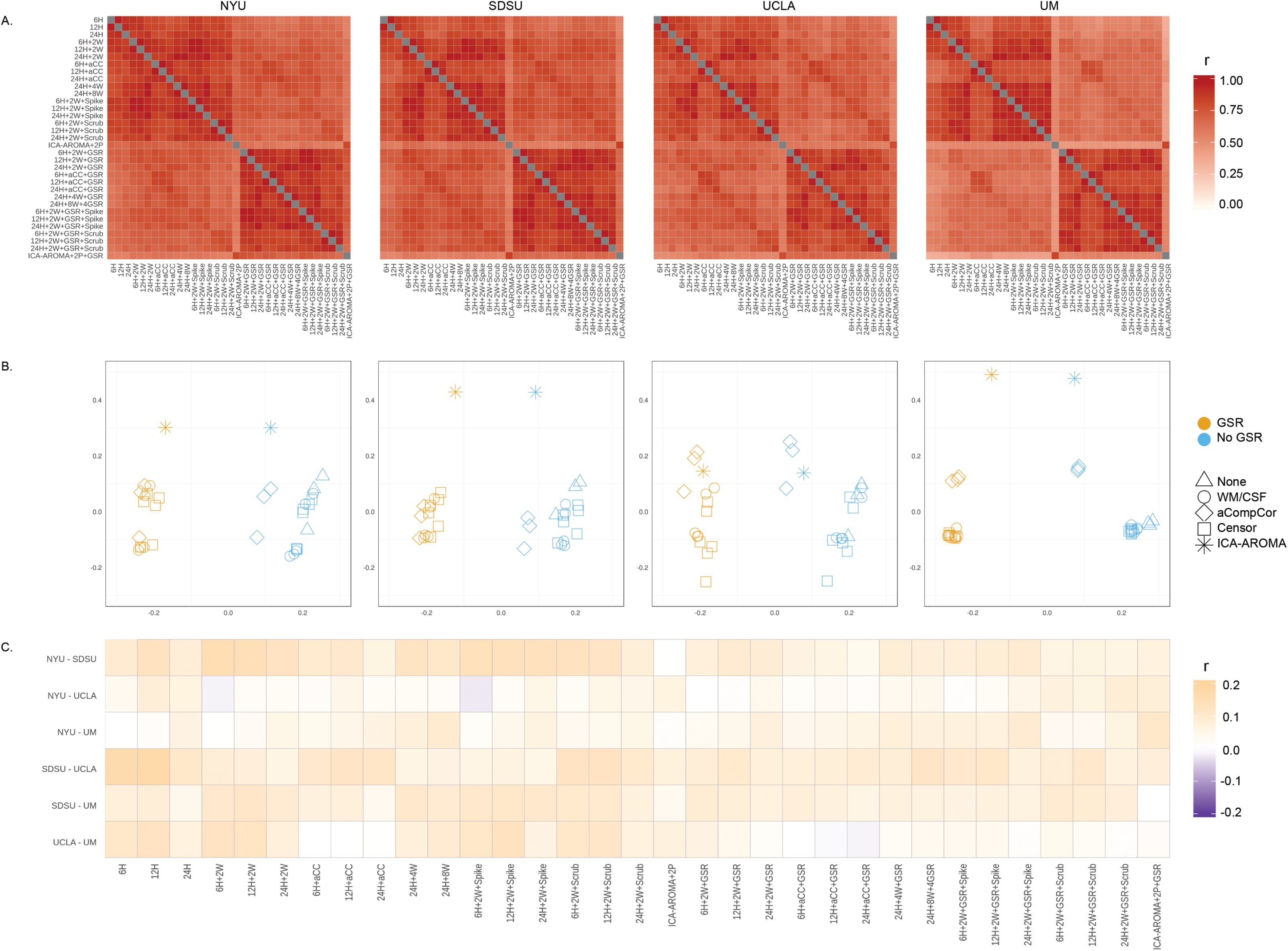
Consistency of group differences in functional connectivity across pipelines and across sites. (A) Spearman’s correlation coefficients between group-difference z-maps across pipelines, and (B) MDS showing relative distance between pipelines, indicate that GSR and ICA-AROMA were different from other strategies. (C) Spearman’s correlation coefficients between group-difference z-maps were consistently low across sites for all pipelines.

The pattern of group differences was not replicable across data sites, regardless of which pipeline was used (Figure 4 and Figure 5). The correlations of group-difference z-maps between sites were consistently low (r = 0.07±0.04, Figure 4C). Pipelines without GSR resulted in slightly higher between-site similarity (mean with GSR: r = 0.061; mean without GSR: r= 0.074; z = 2.22, *p* = 0.03). Even the most different edges between groups within each site rarely overlapped with another site (ASD > control, n = 19.08 ± 11.05; ASD < control, n = 16.66 ± 7.62). Permutation tests indicated that the total number of edges overlapping between two sites was not reliably higher than chance for most pairwise comparisons, with the exception of many pipelines from SDSU-UCLA (*ps* <= 0.05, FDR corrected for six comparisons; note that this analysis did not correct for number of pipelines tested; see Figure 5A). Figure 5B shows the overlap of the 500 most different positive (ASD > control) and negative (control > ASD) edges across the four sites from several representative pipelines. A very small number of edges overlapped between three sites, but these edges varied across pipelines, and no edges overlapped across more than three out of the four sites, in any pipeline.

**Figure 5.**
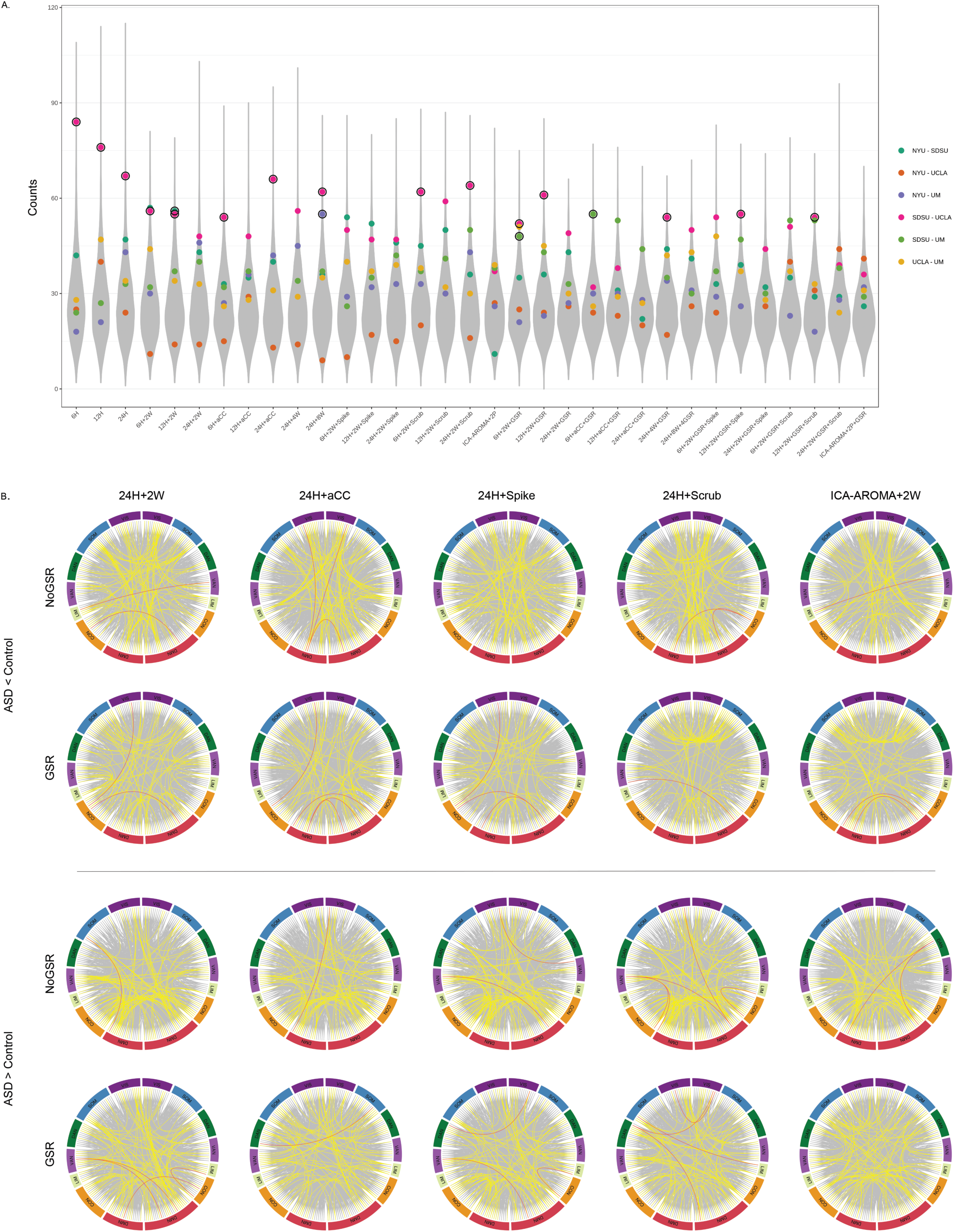
Edge-level overlap between sites. This analysis includes only those edges showing the greatest difference between the ASD and control groups (500 ASD>Control and 500 Control > ASD for each site). (A) The total number of edges that overlapped between sites for each pairwise comparison, represented by a colored dot. The gray distribution is the combined null distribution derived from permutation testing. Dots outlined in black are those identified as significantly higher than chance (q < 0.05, FDR corrected for the six pairwise comparisons within each pipeline). (B) The circular plots show overlapping edges for several different representative pipelines (24H+2W, 24H+aCC, 24H+2W+Spike, 24H+2W+Scrub, 24H+2W+GSR, 24H+aCC+GSR, 24H+2W+GSR+Spike, and 24P+2P+GSR+Scrub, ICA-AROMA+2W, ICA-AROMA+2W+GSR). Line color indicates the number of overlapping sites: grey = 1; yellow = 2; red =3. Note no edge appeared in all sites (four times) across any of the 33 pipelines. VIS: visual network; SOM: somatomotor network; DAN: dorsal attention network; VAN: ventral attention network; LIM: limbic network; CON: control network; DMN: default mode network.

In addition to the fine ROI edge-level resolution, we also examined the consistency of group differences at a larger-scale network level. The pairwise correlation analysis showed the ASD-control group differences at the 17-network level were still inconsistent across data sites (*r* = 0.05±0.27). Figure 6 shows that most of the correlation coefficients between z-maps of each pair of sites were not significant after multiple comparison correction for six pairwise comparisons (no correction for the 33 pipelines to avoid being overly conservative). The between-site similarity varied across pipelines, without significant differences between pipelines with GSR and without GSR (*z* = 0.88; *p* = 0.38).

**Figure 6.**
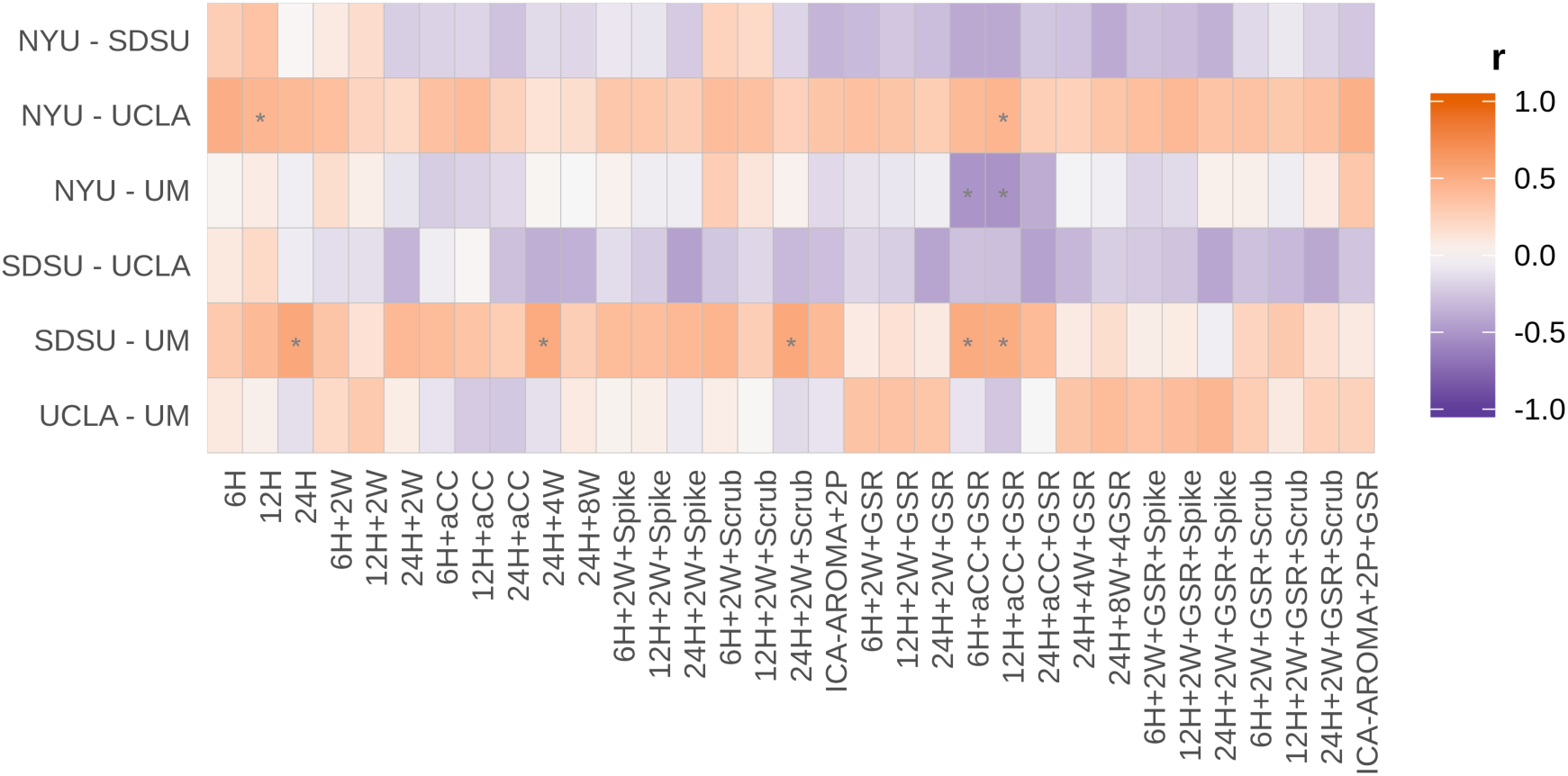
Inconsistency of group differences at the network level across data sites. Note that here, compared to Figure 4C, the results are more variable. \**p* <= 0.05, FDR corrected for six pairwise comparisons. Note also that even where significant differences were identified (e.g., 6H+aCC+GSR), the directionality of the correlation coefficients were inconsistent across pairwise site comparisons (i.e., within columns).

## Discussion

This study examined whether replicable group-level differences between ASD and control groups can be obtained across independently acquired datasets, and how such replicability may vary as a function of preprocessing pipelines. Although basic connectome architecture was highly similar across acquisition sites, regardless of preprocessing pipelines, evidence for replicable group-level ASD-control differences was largely absent. While concerning, it is not altogether surprising as this result is largely consistent with the varied and often conflicting published literature in ASD when taken as a whole – for example, even the basic directionality of effects is still debated (i.e., systematic overconnectivity, underconnectivity, both, or neither).

Here, we show that the lack of replicable ASD-control differences cannot be attributed to the choice of denoising strategy. First, within each site, the pattern of group differences remained largely similar regardless of which denoising strategy was used, as long as the use of GSR was held constant (Figure 3; discussed further below). Second, no particular denoising strategy led to consistently greater across-site replication – i.e., the degree of replication did not improve in any meaningful way with any particular approach (e.g., GSR vs. not). Importantly, this lack of replication was specific to group-level differences and did not extend to basic connectome architecture – when comparing average connectomes across sites, we found a very high degree of similarity, again regardless of denoising procedure. Based on these results, we conclude that while preprocessing may still contribute in part to the lack of replication seen across studies (as it certainly adds variability, and especially with or without GSR), these differences may not be the major factor accounting for such inconsistencies and suggest that other site-level factors play a more significant role.

If differences in denoising strategies cannot adequately explain the lack of across-site replication, an important question is what other factors may account for it. There are at least four possibilities: 1) specific scanner/acquisition/procedural differences; 2) subject-level (cohort) differences; 3) differences in post-processing analysis – e.g., the scale or level (region-of-interest, whole connectome, or network levels); 4) small, hard to detect, or even non-existent differences in functional connectivity in ASD. We unpack these possibilities in the following paragraphs, with each having specific implications for design and analysis of future studies.

On the data collection side, it is possible that uncontrolled factors (including some that remain uncontrolled in the present study) contribute to this lack of replication. These factors include scanner and acquisition parameter differences (e.g., pulse sequence, voxel size, phase encoding directions, scanner manufacturer, etc; Yamashita et al., 2019), as well as experimental procedural differences (e.g., eyes open or closed, experiences immediately preceding the functional scan; Nair et al., 2018). Fortunately, these factors, while they do contribute to across-site variance, tend to be small in terms of effect size (Brown et al., 2011; Dansereau et al., 2017; Noble et al., 2017) or result in localized differences (Nair et al., 2018), consistent with our finding that group-average connectomes were highly reliable across sites. To further increase chances of replication, either *a priori* coordination and standardization of procedures (Glover et al., 2012) or the implementation of post-processing methods designed to increase multisite data harmonization would both be possibilities (Yamashita et al., 2019; Yu et al., 2018).

Another factor related to data collection that potentially underlies our inability to replicate across sites could be subject-level (i.e., cohort) differences or biases (Yamashita et al., 2019). A non-exhaustive list of these factors includes ASD severity, cognitive level, co-morbidities, treatment history and current treatment status (e.g., medication), basic demographic factors including age, sex, race, ethnicity, education, socioeconomic status, and so on. These cohort differences emerge both from practical constraints (e.g., regional biases in terms of participant demographics in different locations) and from the various choices made regarding the recruitment process (e.g., the types of recruitment channels such as clinics vs. communities, and any specific inclusionary and exclusionary criteria). There are several options to remedy these issues. One could apply tightly specified and standardized criteria to match participants across a host of these factors, but in doing so the generalizability of the findings to the broader ASD condition is reduced. A more practical consideration is that attempting to better match sites on some of these factors would result in smaller sample sizes -- for example, in our study, we excluded 184 participants (nearly 31%) from just these four sites in order to better closely match sites on just one of these factors (age). However, it is not necessarily the case that applying more restrictive criteria is always better than including more participants (Abraham et al., 2017). Another way to proceed is to identify the critical factors or grouping of factors that explain significant variance in the data (Smith et al., 2015), and statistically control for those. Other proposals have suggested increasing sampling diversity by collecting relatively small numbers of participants at many different sites, rather than many participants at one site (Dansereau et al., 2017; Yamashita et al., 2019). One recent study (Holiga et al., 2019) that reported replicable findings using the ABIDE dataset combined data across multiple sites as opposed to treating each ABIDE site separately as in the present work – however, effect sizes were smaller in these aggregated samples than in data acquired at a single site. Regardless of the approach one uses, accounting for these subject-level differences is likely an important consideration, as recent work has highlighted that subject-level factors explain more variance than site-level factors (Brown et al., 2011; Dansereau et al., 2017; Gountouna et al., 2010; Noble et al., 2017).

On the analysis side, it is important to note that our findings of a lack of replication are specific to our particular analyses using both whole connectome ROIs-level and a large-scale network-level organization, and do not rule out the possible existence of any other replicable group-level effects in ASD. It is very possible that replicable results could be found when considering the very same data at a different scale or resolution, or with that data analyzed in a different way. For example, King and colleagues (King et al., 2018) found replicable atypical temporal dynamics in rs-fMRI timecourses. Holiga et al. (2019) recently found replicable results regarding functional connectivity in ASD across four very large datasets that also included ABIDE data. However, another recent study by King and colleagues (2019) assessed a number of different measures of functional connectivity in ASD and found weak evidence of generalizability across sites. Other studies have used machine learning approaches to generalize to independently acquired datasets (e.g., Abraham et al., 2017; Yahata et al., 2016). In one of these (Abraham et al., 2017), prediction accuracy was affected by parcellation method, suggesting that replicability may be sensitive to these sorts of analysis choices (e.g., spatial normalization, parcellation; Dadi et al., 2019). Additionally, different scales of connectivity analysis exhibit different sensitivities and vulnerabilities to site effects (Noble et al., 2017), demonstrating a complex and intertwined relationship between many of the factors discussed above. We should mention, however, that although there are different ways of dividing and grouping the data, these approaches mostly still fundamentally rest on the ability to accurately and reliably measure edge-level differences in ASD (e.g., Yahata et al., 2016; see Figure 16a in King et al., 2019). For example, more complex statistical constructs that can be used to compare brain organization between groups (e.g., graph theoretic network measures; (He et al., 2018; Rubinov & Sporns, 2010) fundamentally must build upon reliable and replicable measurement of connectomes. Thus, lack of replication as described in the present work should be of concern to researchers.

The final possibility that ought to be considered is that functional connectivity differences in ASD are very small and difficult or impossible to detect with current technology. While various functional connectivity differences in ASD have been reported in previous studies, the overall lack of consensus is concerning. The growing number of studies that now examine and, in some cases, demonstrate out-of-sample replication provide hope that such replicable signals do in fact exist (Holiga et al., 2019; Yahata et al., 2016). But, because of the above factors and in addition to a host of others (e.g., motion), small differences may be easily obscured (Tyszka, Kennedy, Paul, & Adolphs, 2014). Furthermore, case-control comparisons can easily obscure non-shared, heterogeneous patterns of differences in ASD, and might require different and individually-sensitive analytic approaches (Byrge et al., 2015; Dubois & Adolphs, 2016; Marquand, Rezek, Buitelaar, & Beckmann, 2016; Marquand et al., 2019). How to reliably detect these differences by using current neuroimaging methodologies and analytic approaches remains an open question for future work.

What does this all mean? The pessimistic view would be that researchers should give up on searching for common group-level effects in ASD. However, we believe that this conclusion would be very premature for a number of reasons. (1) It is possible that effects are heterogeneous across participants, so group-level analysis starting with the assumption of homogeneous groups may be both largely underpowered and not able to fully account for the group level variance. (2) It is possible that improvements in detecting signal in the face of the large amounts of measurement noise that plague resting-state analyses will eventually unmask important group-level differences. In this case, if it is a detection problem, continued advances in acquisition and analysis methodology may get us closer to detecting reliable differences in ASD. (3) Additional experimental procedures can be employed to ensure more reliable estimates of an individual’s connectome. For example, collecting more data from each individual participant can reduce measurement noise and ensure greater confidence in the results via within-sample replication (Anderson, King, & Anderson, 2019; Byrge & Kennedy, 2019; Finn et al., 2015; King et al., 2019; Nee, 2019), prior to attempting across-site replication.

While our results suggest that lack of replication cannot be solely attributed to differences in denoising procedures (since using the same preprocessing procedures did not increase across-site replication), this does not mean that they are entirely inconsequential. Here, we show that, while there are essentially an unconstrained number of choices for preprocessing, some of these choices have a more significant impact on the results than others (though not necessarily in a consistent way). Figure 3 demonstrates that one of the most significant factors is whether or not GSR is included as a preprocessing step. Its inclusion resulted in slightly more similar group-averaged connectomes across sites -- however, whether more similar group-averaged connectomes is a good thing or not remains unclear. The positive interpretation of this finding is that GSR helps to eliminate measurement noise (Power et al., 2014; Power et al., 2017; Byrge & Kennedy, 2018; Ciric et al., 2017; Parkes et al., 2018), resulting in more similar connectomes, whereas the less positive interpretation is that GSR eliminates individual variation that might be of interest or distorts group-level differences (Gotts et al., 2013; Scholvinck et al., 2010; Uddin, 2017; Yang et al., 2014). Our results cannot disambiguate these possibilities from one another. Furthermore, in terms of group differences, we found that the effects of GSR on across-site replicability were not consistent, and instead depended on which specific sites were compared to one another (see Figure 4, middle panel, and Figure 6). For some site comparisons, use of GSR increased similarity between them, whereas for others it decreased it, and yet others where it was unchanged, suggesting a complex interaction between the use of GSR and site-level factors.

In addition to the possible factors already discussed above that may limit the detection of reliable group effects, some additional limitations of this study are worth mentioning. One criticism is that correlations between whole connectome group difference z-maps are perhaps a relatively insensitive way to examine this data. For instance, a localized difference in a small number of edges or nodes would easily be obscured in the present whole-brain analyses. However, we did also examine only the edges that differed most between groups, and also examined data aggregated at the network level – both yielded poor replicability of results. Another limitation of the present study is the relatively small sample sizes. This was a consequence of both carefully matching groups by age and also applying strict quality control (i.e., movement thresholds, anatomical image quality requirements). However, we note that our sample size was sufficiently powered to detect medium-large to large effects within each dataset, suggesting that possible replicable differences must be smaller than this. As shown in the literature survey (Supplementary Figure S1), although there may be a recent growing trend to use larger datasets (primarily aggregated from ABIDE), many studies still use single site data with limited sample size. Indeed, the median sample sizes in these published studies (ASD: n = 35; TD: n = 38) are approximately equal to or smaller than the four sample sizes used in the present work – studies that form the basis of our understanding of functional connectivity abnormalities in ASD. Another limitation is that the present study included a relatively large age range from 10-20 years, corresponding to a broad neurodevelopmental period spanning childhood through adolescence and into young adulthood. This age range is not uncommon among previous studies, as showed in the literature survey (Supplementary Figure S1). It is possible that more consistent effects would be identified if the age was constrained even further – however, further restricting the range would have reduced the number of sites and subjects that we could have included.

In sum, the present study demonstrated that the choice of denoising pipeline is not the main factor underlying the lack of replication of group differences in ASD. Instead, the most parsimonious explanation is that group-level differences are small or non-existent, and/or swamped by site and sample effects. However, we remain optimistic that continued developments toward improving methodology and approaches will help to eventually reveal reliable patterns of functional connectivity alterations in ASD. These results highlight the need to continue examining reliability of findings going forward, and demonstrate that approaches that improve sensitivity to detect disorder-related alterations are still needed.

## Acknowledgement

This work was supported by the NIH (R01MH110630 and R00MH094409 to DPK), and NICHD (T32HD007475 Postdoctoral Traineeship to LB). The authors gratefully acknowledge the contributors and organizers of the Autism Brain Imaging Data Exchange, as well as Dr. Linden Parkes and colleagues for making their preprocessing code publicly available. ABIDE I is supported by NIMH (K23MH087770; R03MH096321), the Leon Levy Foundation, Joseph P. Healy, and the Stavros Niarchos Foundation. ABIDE II is supported by NIMH (5R21MH107045), Nathan S. Kline Institute of Psychiatric Research, Joseph P. Healey, Phyllis Green, and Randolph Cowen. We also acknowledge the Indiana University Pervasive Technology Institute for providing HPC (Big Red II, Karst, Carbonate), visualization, and storage resources, which were supported in part by Lilly Endowment, Inc., the Indiana METACyt Initiative, and based upon work supported by the National Science Foundation under Grant No. CNS-0521433.

## Declarations of interest

none

## Notes

http://fcon_1000.projects.nitrc.org/indi/abide/

## References

Abraham, A., Milham, M. P., Di Martino, A., Craddock, R. C., Samaras, D., Thirion, B., & Varoquaux, G. (2017). Deriving reproducible biomarkers from multi-site resting-state data: An Autism-based example. Neuroimage, 147, 736–745. doi:10.1016/j.neuroimage.2016.10.045

Anderson, A. N., King, J. B., & Anderson, J. S. (2019). Neuroimaging in Psychiatry and Neurodevelopment: why the emperor has no clothes. The British Journal of Radiology, 20180910. doi:10.1259/bjr.20180910

Avants, B. B., Epstein, C. L., Grossman, M., & Gee, J. C. (2008). Symmetric diffeomorphic image registration with cross-correlation: evaluating automated labeling of elderly and neurodegenerative brain. Med Image Anal, 12(1), 26–41. doi:10.1016/j.media.2007.06.004

Benjamini, Y., & Hochberg, Y. (1995). Controlling the False Discovery Rate - a Practical and Powerful Approach to Multiple Testing. Journal of the Royal Statistical Society Series B-Statistical Methodology, 57(1), 289–300.

Birn, R. M. (2012). The role of physiological noise in resting-state functional connectivity. Neuroimage, 62(2), 864–870. doi:10.1016/j.neuroimage.2012.01.016

Biswal, B., Yetkin, F. Z., Haughton, V. M., & Hyde, J. S. (1995). Functional Connectivity in the Motor Cortex of Resting Human Brain Using Echo-Planar Mri. Magnetic Resonance in Medicine, 34(4), 537–541. doi: DOI 10.1002/mrm.1910340409

Brown, G. G., Mathalon, D. H., Stern, H., Ford, J., Mueller, B., Greve, D. N., … Function Biomedical Informatics Research, N. (2011). Multisite reliability of cognitive BOLD data. Neuroimage, 54(3), 2163–2175. doi:10.1016/j.neuroimage.2010.09.076

Byrge, L., Dubois, J., Tyszka, J. M., Adolphs, R., & Kennedy, D. P. (2015). Idiosyncratic brain activation patterns are associated with poor social comprehension in autism. J Neurosci, 35(14), 5837–5850. doi:10.1523/JNEUROSCI.5182-14.2015

Byrge, L., & Kennedy, D. P. (2018). Identifying and characterizing systematic temporally-lagged BOLD artifacts. Neuroimage, 171, 376–392. doi:10.1016/j.neuroimage.2017.12.082

Byrge, L., & Kennedy, D. P. (2019). High-accuracy individual identification using a “thin slice” of the functional connectome. Network Neuroscience, 3(2), 363–383. doi:10.1162/netn_a_00068

Ciric, R., Wolf, D. H., Power, J. D., Roalf, D. R., Baum, G. L., Ruparel, K., Satterthwaite, T. D. (2017). Benchmarking of participant-level confound regression strategies for the control of motion artifact in studies of functional connectivity. Neuroimage, 154, 174–187. doi:10.1016/j.neuroimage.2017.03.020

Dadi, K., Rahim, M., Abraham, A., Chyzhyk, D., Milham, M., Thirion, B., & Varoquaux, G. (2019). Benchmarking functional connectome-based predictive models for resting-state fMRI. Neuroimage, 192, 115–134. doi: https://doi.org/10.1016/j.neuroimage.2019.02.062

Dansereau, C., Benhajali, Y., Risterucci, C., Pich, E. M., Orban, P., Arnold, D., & Bellec, P. (2017). Statistical power and prediction accuracy in multisite resting-state fMRI connectivity. Neuroimage, 149, 220–232. doi:10.1016/j.neuroimage.2017.01.072

Di Martino, A., O’Connor, D., Chen, B., Alaerts, K., Anderson, J. S., Assaf, M., … Milham, M. P. (2017). Data Descriptor: Enhancing studies of the connectome in autism using the autism brain imaging data exchange II. Scientific Data, 4. doi:10.1038/sdata.2017.10

Di Martino, A., Yan, C. G., Li, Q., Denio, E., Castellanos, F. X., Alaerts, K., … Milham, M. P. (2014). The autism brain imaging data exchange: towards a large-scale evaluation of the intrinsic brain architecture in autism. Mol Psychiatry, 19(6), 659–667. doi:10.1038/mp.2013.78

Dubois, J., & Adolphs, R. (2016). Building a Science of Individual Differences from fMRI. Trends Cogn Sci, 20(6), 425–443. doi:10.1016/j.tics.2016.03.014

Finn, E. S., Shen, X. L., Scheinost, D., Rosenberg, M. D., Huang, J., Chun, M. M., … Constable, R. T. (2015). Functional connectome fingerprinting: identifying individuals using patterns of brain connectivity. Nature Neuroscience, 18(11), 1664–1671. doi:10.1038/nn.4135

Floris, D. L., Lai, M.-C., Nath, T., Milham, M. P., & Di Martino, A. (2018). Network-specific sex differentiation of intrinsic brain function in males with autism. Molecular Autism, 9(1), 17. doi:10.1186/s13229-018-0192-x

Glover, G. H., Mueller, B. A., Turner, J. A., van Erp, T. G., Liu, T. T., Greve, D. N., … Potkin, S. G. (2012). Function biomedical informatics research network recommendations for prospective multicenter functional MRI studies. J Magn Reson Imaging, 36(1), 39–54. doi:10.1002/jmri.23572

Gotts, S. J., Saad, Z. S., Jo, H. J., Wallace, G. L., Cox, R. W., & Martin, A. (2013). The perils of global signal regression for group comparisons: a case study of Autism Spectrum Disorders. Front Hum Neurosci, 7, 356. doi:10.3389/fnhum.2013.00356

Gountouna, V. E., Job, D. E., McIntosh, A. M., Moorhead, T. W., Lymer, G. K., Whalley, H. C., … Lawrie, S. M. (2010). Functional Magnetic Resonance Imaging (fMRI) reproducibility and variance components across visits and scanning sites with a finger tapping task. Neuroimage, 49(1), 552–560. doi:10.1016/j.neuroimage.2009.07.026

Greicius, M. D., Krasnow, B., Reiss, A. L., & Menon, V. (2003). Functional connectivity in the resting brain: a network analysis of the default mode hypothesis. Proc Natl Acad Sci U S A, 100(1), 253–258. doi:10.1073/pnas.0135058100

Hahamy, A., Behrmann, M., & Malach, R. (2015). The idiosyncratic brain: distortion of spontaneous connectivity patterns in autism spectrum disorder. Nature Neuroscience, 18(2), 302–309. doi:10.1038/nn.3919

He, Y., Lim, S., Fortunato, S., Sporns, O., Zhang, L., Qiu, J., … Zuo, X. N. (2018). Reconfiguration of Cortical Networks in MDD Uncovered by Multiscale Community Detection with fMRI. Cerebral Cortex, 28(4), 1383–1395. doi:10.1093/cercor/bhx335

Holiga, S., Hipp, J. F., Chatham, C. H., Garces, P., Spooren, W., D’Ardhuy, X. L., … Dukart, J. (2019). Patients with autism spectrum disorders display reproducible functional connectivity alterations. Science Translational Medicine, 11(481). doi:10.1126/scitranslmed.aat9223

Hull, J. V., Dokovna, L. B., Jacokes, Z. J., Torgerson, C. M., Irimia, A., & Van Horn, J. D. (2016). Resting-State Functional Connectivity in Autism Spectrum Disorders: A Review. Front Psychiatry, 7, 205. doi:10.3389/fpsyt.2016.00205

Jenkinson, M., Bannister, P., Brady, M., & Smith, S. (2002). Improved optimization for the robust and accurate linear registration and motion correction of brain images. Neuroimage, 17(2), 825–841. doi:10.1006/nimg.2002.1132

Jones, T. B., Bandettini, P. A., Kenworthy, L., Case, L. K., Milleville, S. C., Martin, A., & Birn, R. M. (2010). Sources of group differences in functional connectivity: an investigation applied to autism spectrum disorder. Neuroimage, 49(1), 401–414. doi:10.1016/j.neuroimage.2009.07.051

King, J. B., Prigge, M. B. D., King, C. K., Morgan, J., Dean, D. C., Freeman, A., … Anderson, J. S. (2018). Evaluation of Differences in Temporal Synchrony Between Brain Regions in Individuals With Autism and Typical Development. Jama Network Open, 1(7). doi:10.1001/jamanetworkopen.2018.4777

King, J. B., Prigge, M. B. D., King, C. K., Morgan, J., Weathersby, F., Fox, J. C., … Anderson, J. S. (2019). Generalizability and reproducibility of functional connectivity in autism. Mol Autism, 10, 27. doi:10.1186/s13229-019-0273-5

Lemieux, L., Salek-Haddadi, A., Lund, T. E., Laufs, H., & Carmichael, D. (2007). Modelling large motion events in fMRI studies of patients with epilepsy. Magn Reson Imaging, 25(6), 894–901. doi:10.1016/j.mri.2007.03.009

Marquand, A. F., Rezek, I., Buitelaar, J., & Beckmann, C. F. (2016). Understanding Heterogeneity in Clinical Cohorts Using Normative Models: Beyond Case-Control Studies. Biological Psychiatry, 80(7), 552–561. doi:10.1016/j.biopsych.2015.12.023

Marquand, A. F., Kia, S. M., Zabihi, M., Wolfers, T., Buitelaar, J. K., & Beckmann, C. F. (2019). Conceptualizing mental disorders as deviations from normative functioning. Molecular Psychiatry. doi:10.1038/s41380-019-0441-1

Müller, R.-A., Shih, P., Keehn, B., Deyoe, J. R., Leyden, K. M., & Shukla, D. K. (2011). Underconnected, but How? A Survey of Functional Connectivity MRI Studies in Autism Spectrum Disorders. Cerebral Cortex, 21(10), 2233–2243. doi:10.1093/cercor/bhq296

Muschelli, J., Nebel, M. B., Caffo, B. S., Barber, A. D., Pekar, J. J., & Mostofsky, S. H. (2014). Reduction of motion-related artifacts in resting state fMRI using aCompCor. Neuroimage, 96, 22–35. doi:10.1016/j.neuroimage.2014.03.028

Nair, S., Jao Keehn, R. J., Berkebile, M. M., Maximo, J. O., Witkowska, N., & Muller, R. A. (2018). Local resting state functional connectivity in autism: site and cohort variability and the effect of eye status. Brain Imaging Behav, 12(1), 168–179. doi:10.1007/s11682-017-9678-y

Nee, D. E. (2019). fMRI replicability depends upon sufficient individual-level data. Communications Biology, 2(1), 130. doi:10.1038/s42003-019-0378-6

Noble, S., Scheinost, D., Finn, E. S., Shen, X., Papademetris, X., McEwen, S. C., … Constable, R. T. (2017). Multisite reliability of MR-based functional connectivity. Neuroimage, 146, 959–970. doi:10.1016/j.neuroimage.2016.10.020

Parkes, L., Fulcher, B., Yucel, M., & Fornito, A. (2018). An evaluation of the efficacy, reliability, and sensitivity of motion correction strategies for resting-state functional MRI. Neuroimage, 171, 415–436. doi:10.1016/j.neuroimage.2017.12.073

Power, J. D., Barnes, K. A., Snyder, A. Z., Schlaggar, B. L., & Petersen, S. E. (2012). Spurious but systematic correlations in functional connectivity MRI networks arise from subject motion. Neuroimage, 59(3), 2142–2154. doi:10.1016/j.neuroimage.2011.10.018

Power, J. D., Mitra, A., Laumann, T. O., Snyder, A. Z., Schlaggar, B. L., & Petersen, S. E. (2014). Methods to detect, characterize, and remove motion artifact in resting state fMRI. Neuroimage, 84, 320–341. doi:10.1016/j.neuroimage.2013.08.048

Power, J. D., Plitt, M., Laumann, T. O., & Martin, A. (2017). Sources and implications of whole-brain fMRI signals in humans. Neuroimage, 146, 609–625. doi:10.1016/j.neuroimage.2016.09.038

Pruim, R. H. R., Mennes, M., van Rooij, D., Llera, A., Buitelaar, J. K., & Beckmann, C. F. (2015). ICA-AROMA: A robust ICA-based strategy for removing motion artifacts from fMRI data. Neuroimage, 112, 267–277. doi:10.1016/j.neuroimage.2015.02.064

Pua, E. P. K., Malpas, C. B., Bowden, S. C., & Seal, M. L. (2018). Different brain networks underlying intelligence in autism spectrum disorders. Human Brain Mapping, 39(8), 3253–3262. doi:10.1002/hbm.24074

Rubinov, M., & Sporns, O. (2010). Complex network measures of brain connectivity: uses and interpretations. Neuroimage, 52(3), 1059–1069. doi:10.1016/j.neuroimage.2009.10.003

Satterthwaite, T. D., Elliott, M. A., Gerraty, R. T., Ruparel, K., … Loughead, J., Calkins, M. E., … Wolf, D. H. (2013). An improved framework for confound regression and filtering for control of motion artifact in the preprocessing of resting-state functional connectivity data. Neuroimage, 64, 240–256. doi:10.1016/j.neuroimage.2012.08.052

Satterthwaite, T. D., Wolf, D. H., Loughead, J., Ruparel, K., Elliott, M. A., Hakonarson, H., … Gur, R. E. (2012). Impact of in-scanner head motion on multiple measures of functional connectivity: relevance for studies of neurodevelopment in youth. Neuroimage, 60(1), 623–632. doi:10.1016/j.neuroimage.2011.12.063

Schaefer, A., Kong, R., Gordon, E. M., Laumann, T. O., Zuo, X. N., Holmes, A. J., … Thomas, B. T. (2018). Local-Global Parcellation of the Human Cerebral Cortex from Intrinsic Functional Connectivity MRI. Cerebral Cortex, 28(9), 3095–3114. doi:10.1093/cercor/bhx179

Scholvinck, M. L., Maier, A., Ye, F. Q., Duyn, J. H., & Leopold, D. A. (2010). Neural basis of global resting-state fMRI activity. Proc Natl Acad Sci U S A, 107(22), 10238–10243. doi:10.1073/pnas.0913110107

Smith, S. M., Jenkinson, M., Woolrich, M. W., Beckmann, C. F., Behrens, T. E. J., Johansen-Berg, H., … Matthews, P. M. (2004). Advances in functional and structural MR image analysis and implementation as FSL. Neuroimage, 23, S208–S219. doi:10.1016/j.neuroimage.2004.07.051

Smith, S. M., Nichols, T. E., Vidaurre, D., Winkler, A. M., Behrens, T. E., Glasser, M. F., … Miller, K. L. (2015). A positive-negative mode of population covariation links brain connectivity, demographics and behavior. Nature Neuroscience, 18(11), 1565–1567. doi:10.1038/nn.4125

Turner, J. A., Damaraju, E., van Erp, T. G., Mathalon, D. H., Ford, J. M., Voyvodic, J., … Calhoun, V. D. (2013). A multi-site resting state fMRI study on the amplitude of low frequency fluctuations in schizophrenia. Front Neurosci, 7, 137. doi:10.3389/fnins.2013.00137

Tyszka, J. M., Kennedy, D. P., Paul, L. K., & Adolphs, R. (2014). Largely typical patterns of resting-state functional connectivity in high-functioning adults with autism. Cerebral Cortex, 24(7), 1894–1905. doi:10.1093/cercor/bht040

Uddin, L. Q. (2017). Mixed Signals: On Separating Brain Signal from Noise. Trends Cogn Sci, 21(6), 405–406. doi:10.1016/j.tics.2017.04.002

Van Dijk, K. R., Sabuncu, M. R., & Buckner, R. L. (2012). The influence of head motion on intrinsic functional connectivity MRI. Neuroimage, 59(1), 431–438. doi:10.1016/j.neuroimage.2011.07.044

Yahata, N., Morimoto, J., Hashimoto, R., Lisi, G., Shibata, K., Kawakubo, Y., … Kawato, M. (2016). A small number of abnormal brain connections predicts adult autism spectrum disorder. Nat Commun, 7, 11254. doi:10.1038/ncomms11254

Yamashita, A., Yahata, N., Itahashi, T., Lisi, G., Yamada, T., Ichikawa, N., … Imamizu, H. (2019). Harmonization of resting-state functional MRI data across multiple imaging sites via the separation of site differences into sampling bias and measurement bias. PLOS Biology, 17(4), e3000042. doi:10.1371/journal.pbio.3000042

Yan, C. G., Cheung, B., Kelly, C., Colcombe, S., Craddock, R. C., Di Martino, A., … Milham, M. P. (2013). A comprehensive assessment of regional variation in the impact of head micromovements on functional connectomics. Neuroimage, 76, 183–201. doi:10.1016/j.neuroimage.2013.03.004

Yan, C. G., Craddock, R. C., Zuo, X. N., Zang, Y. F., & Milham, M. P. (2013). Standardizing the intrinsic brain: Towards robust measurement of inter-individual variation in 1000 functional connectomes. Neuroimage, 80, 246–262. doi:10.1016/j.neuroimage.2013.04.081

Yang, G. J., Murray, J. D., Repovs, G., Cole, M. W., Savic, A., Glasser, M. F., … Anticevic, A. (2014). Altered global brain signal in schizophrenia. Proc Natl Acad Sci U S A, 111(20), 7438–7443. doi:10.1073/pnas.1405289111

Yeo, B. T., Krienen, F. M., Sepulcre, J., Sabuncu, M. R., Lashkari, D., Hollinshead, M., … Buckner, R. L. (2011). The organization of the human cerebral cortex estimated by intrinsic functional connectivity. J Neurophysiol, 106(3), 1125–1165. doi:10.1152/jn.00338.2011

Yu, M., Linn, K. A., Cook, P. A., Phillips, M. L., McInnis, M., Fava, M., … Sheline, Y. I. (2018). Statistical harmonization corrects site effects in functional connectivity measurements from multi-site fMRI data. Hum Brain Mapp, 39(11), 4213–4227. doi:10.1002/hbm.24241

