## Supplementary material for "Non-replication of functional connectivity differences in autism spectrum disorder across multiple sites and denoising strategies": Figure S1

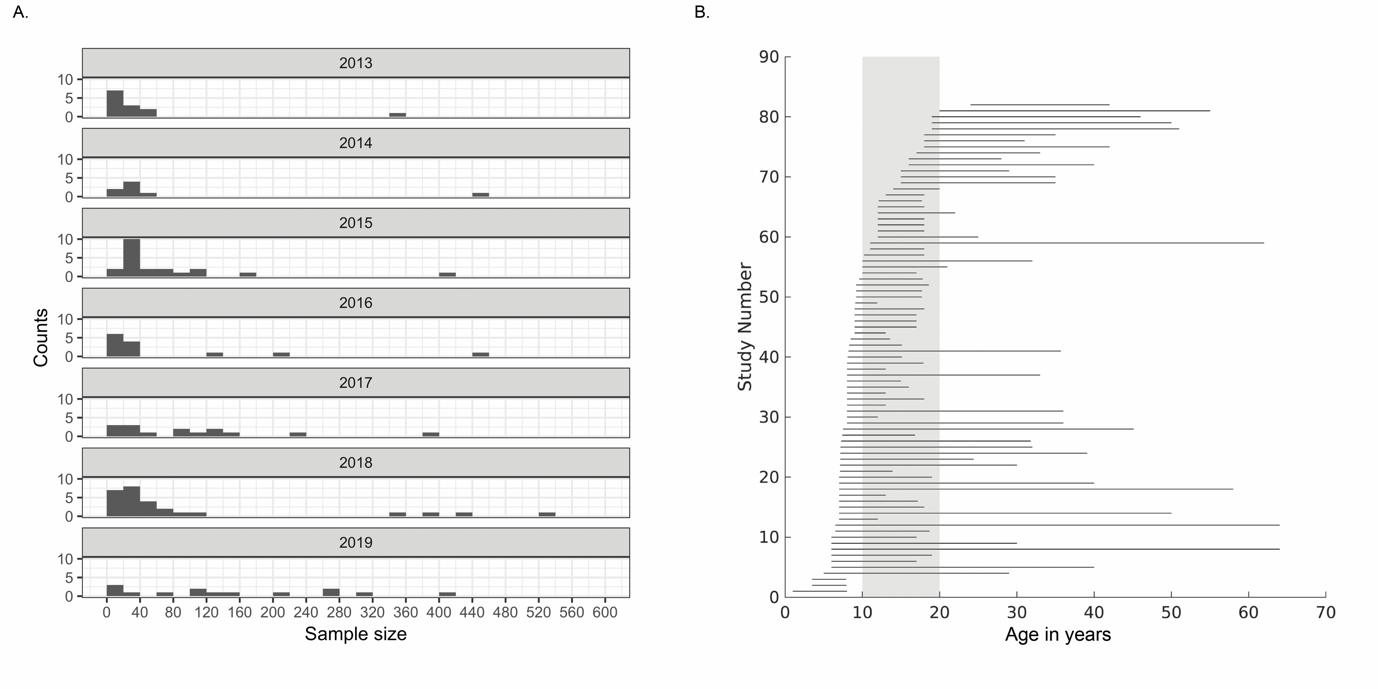

Figure S1. Characteristics of samples used in previous relevant resting-state fMRI studies of ASD from January 2013 to June 2019. A) A histogram of sample sizes of the ASD group across studies, by publication year. B) Age ranges from previous studies (among those that reported it). Each line represents the range for one study; the gray shadowing highlights the range of ages (10-20 years) included in the present study. Note that a large number of published studies include ages spanning this developmental period.
